# The neurotranscriptome of the *Aedes aegypti* mosquito

**DOI:** 10.1101/026823

**Authors:** Benjamin J. Matthews, Carolyn S. McBride, Matthew DeGennaro, Orion Despo, Leslie B. Vosshall

**Affiliations:** Laboratory of Neurogenetics and Behavior, The Rockefeller University, New York, NY 10065 USA; Howard Hughes Medical Institute, New York, NY 10065 USA

**Keywords:** Mosquito, *Aedes aegypti*, mRNA-sequencing, *de novo* genome assembly, host-seeking behavior, neural genes, chemosensory receptors, ion channels, G protein-coupled receptors, gonotrophic cycle, neurogenetics

## Abstract

**Background:** A complete genome sequence and the advent of genome editing open up non-traditional model organisms to mechanistic genetic studies. The mosquito *Aedes aegypti* is an important vector of infectious diseases such as dengue, chikungunya, and yellow fever, and has a large and complex genome, which has slowed annotation efforts. We used comprehensive transcriptomic analysis of adult gene expression to improve the genome annotation and to provide a detailed tissue-specific catalogue of neural gene expression at different adult behavioral states.

**Results:** We carried out deep RNA sequencing across all major peripheral male and female sensory tissues, the brain, and (female) ovary. Furthermore, we examined gene expression across three important phases of the female reproductive cycle, a remarkable example of behavioral switching in which a female mosquito alternates between obtaining blood-meals from humans and laying eggs. Using genome-guided alignments and *de novo* transcriptome assembly, our re-annotation includes 572 new putative protein-coding genes and updates to 13.5% and 50.3% of existing transcripts within coding sequences and untranslated regions, respectively. Using this updated annotation, we detail gene expression in each tissue, identifying large numbers of transcripts regulated by blood-feeding and sexually dimorphic transcripts that may provide clues to the biology of male- and female-specific behaviors, such as mating and blood-feeding, which are areas of intensive study for those interested in vector control.

**Conclusions:** This neurotranscriptome forms a strong foundation for the study of genes in the mosquito nervous system and investigation of sensory-driven behaviors and their regulation. Furthermore, understanding the molecular genetic basis of mosquito chemosensory behavior has important implications for vector control.

## Background

Studies in classic genetic model organisms including the mouse, zebrafish, fly, worm, and yeast have led to major advances in biology. All of these systems have in common a sequenced genome and the ability to carry out forward and reverse genetic manipulations. Non-model organisms, such as the mosquitoes we study, have not been accessible to mechanistic genetic studies until recently. The availability of genomes, next-generation sequencing, and genome editing technologies now make it possible to apply modern genetics to study animals with important and interesting biology previously inaccessible to molecular genetics.

*Aedes aegypti* is the primary vector for dengue, chikungunya, and yellow fever – debilitating diseases that together are responsible for hundreds of millions of infections and thousands of deaths annually worldwide [1]. Female mosquitoes exhibit remarkable behavioral shifts throughout their adult life. *Ae. aegypti* are generally anautogenous, meaning that they do not produce eggs without a blood-meal [2]. Female *Ae. aegypti* use a variety of chemical and physical cues to locate hosts in their environment and to discriminate humans from non-human animals [3–8]. Although male *Ae. aegypti* do not feed on blood, they also respond to host chemosensory cues, perhaps to locate females congregating near humans [9]. At short range, the male locates a potential mate using the specific frequencies generated by a female’s wing-beat [10].

After successfully obtaining a blood-meal, female mosquitoes repress host-seeking behavior [11, 12], and utilize the nutrients in the blood-meal to develop a batch of eggs. A female who has reached this physiological state is known as “gravid”. It is known that egg maturation and the beginning of egg-laying behavior occur between 48 and 96 hours after a blood-meal [11]. Once the eggs have matured, a gravid female uses cues such as humidity and the presence and quality of liquid water to identify a suitable place to lay her eggs, a behavior also known as oviposition [13]. Following oviposition, a female mosquito recovers her attraction to hosts and seeks out new blood-meals to produce successive batches of eggs. This process, including host-seeking, egg maturation, and oviposition, is known as the gonotrophic cycle [11]. Disease transmission by mosquitoes is driven by this cyclical nature of female biting behavior, as a mosquito must first bite an infected host before becoming competent to spread infection to subsequent hosts.

Genetic resources, such as those that have long existed for conventional model organisms, would greatly facilitate investigation into the mechanistic basis of behavior in mosquitoes. While there has been impressive progress in mosquito transgenesis and mutagenesis in the past 20 years [3, 8, 12, 14–22], the large size of the *Ae. aegypti* genome (∼1.3 Gb) and large transposable element load (∼47%) present formidable challenges to genome assembly, physical mapping, and annotation [23–26]. Despite the limitations imposed by incomplete annotation of new mosquito genomes, several studies have profiled gene expression in individual sensory organs in the mosquitoes *Ae. aegypti* [27–29], *Culex quinquefasciatus* [30], *Anopheles gambiae* [31–34], and *Toxorhynchites amboinensis* [34].

Our work builds on these efforts by incorporating biological replicates sequenced at greater depth and from many isolated tissues in parallel in both females at several behavioral states, and in males. This large dataset makes it possible to detect genes expressed at low levels or expressed in only a few neurons, and to identify differential gene expression with statistical confidence. Since the anatomical substrate of host-seeking, egg-laying, and other mosquito behaviors is likely to be distributed across several tissues, parallel transcriptional profiling of multiple tissues in a single study increases the likelihood of capturing the full repertoire of genes involved in these complex behaviors.

To generate a transcriptome of peripheral and central neural tissues (or “neurotranscriptome”) in *Ae. aegypti*, we performed Illumina mRNA-sequencing (RNA-seq) on RNA isolated from male and female tissues. Tissues sampled included the brain, antenna, maxillary palp, proboscis, abdominal tip, legs, and female ovary. To understand the influence of blood-feeding state on gene expression, we performed RNA-seq on a subset of tissues in female mosquitoes at three time-points: prior to a blood-meal (non-blood-fed), at 48 hours following a blood-meal (blood-fed), and at 96 hours following a blood-meal (gravid).

This project, as part of the NIAID VectorBase Driving Biological Projects Initiative [35], set out to accomplish three major goals: 1) to improve the existing annotation of protein-coding genes in the *Ae. aegypti* genome and identify genes not found in the current genome assembly; 2) to catalogue gene expression at the resolution of single tissues in host-seeking female and male mosquitoes; and 3) to identify changes in gene expression that are correlated with blood-feeding state and its associated behavioral changes. This neurotranscriptome significantly enhances the *Ae. aegypti* genome annotation, and identifies a large number of genes whose expression is sexually dimorphic and/or variable across the female gonotrophic cycle. We anticipate that these data will drive studies of the genetic and neural circuit basis of host-seeking and egg-laying behavior in *Ae. aegypti.*

## Results

We profiled *Ae. aegypti* gene expression in tissues of males, and females at three points in their gonotrophic cycle (Figure 1A). To confirm the distinct behavioral states associated with these time points in the mosquitoes used for our RNA-seq study, we measured responses to live hosts and carbon dioxide, and monitored female egg-laying. A uniport olfactometer was used to measure female mosquito attraction to a human forearm, which was presented along with carbon dioxide (CO_2_) to activate the mosquitoes [12]. Non-blood-fed females showed strong attraction to these host cues, while blood-fed females did not respond and gravid females showed only modest host attraction (Figure 1B). Because oviposition is known to release host-seeking suppression [11], egg-laying was prevented prior to behavioral testing or tissue dissection by depriving females of access to water. We confirmed that oviposition behavior was normal in these animals by placing individual blood-fed and gravid female mosquitoes into oviposition vials and scoring the number of eggs laid over an 8 hour period. As expected, blood-fed mosquitoes laid no eggs, but nearly all gravid mosquitoes did (Figure 1C).

**Figure 1.**
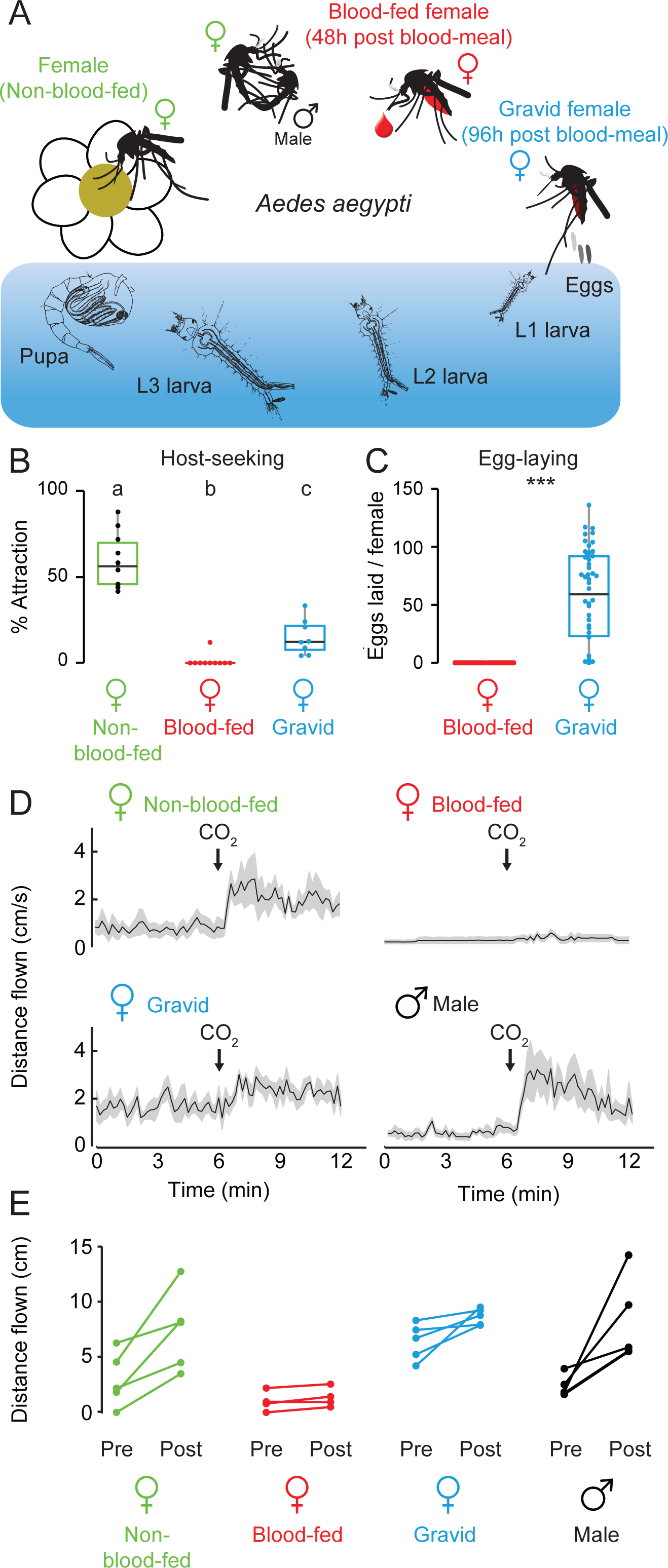
Ae. aegypti life cycle and adult behaviors. (**A**) Schematic of the mosquito life cycle and the female gonotrophic cycle. (**B**) Female host-seeking behavior at three stages of the gonotrophic cycle as assayed in a uniport olfactometer [12] (ANOVA followed by Tukey’s HSD, all differences p < 0.05, n = 8-10 per condition). (**C**) Eggs laid in 8 hours after females were offered an oviposition substrate at 48 or 96 hours after a blood-meal (Welch two-sample t-test, p < 0.001; n = 50 females). (**D**) Response to CO_2_ assayed in a three dimensional flight tracking arena [8]. Mean indicated with the black line and S.E.M. by the gray shading; n=4-5 per condition. (**E**) Summary of flight tracking data, with distance flown quantified in 6-minute periods prior to and following a 40 second pulse of CO_2_. Post-CO_2_ activity of blood-fed females is reduced as compared to all other conditions (ANOVA followed by Tukey’s HSD; p<0.05), and pre-CO_2_ activity of gravid females is increased as compared to all other conditions (ANOVA followed by Tukey’s HSD; p<0.05).

To determine whether the diminished responses to human hosts following blood feeding can be solely attributed to a reduction in sensitivity to CO_2_, we utilized a multi-insect three-dimensional flight-tracking system [8] to assess the response of a group of 20 female mosquitoes to a 40 second pulse of CO_2_ (Figure 1D and 1E). As previously described, non-blood-fed female mosquitoes displayed a robust increase in flight activity in response to CO_2_ [8], while blood-fed females showed no increase in activity following administration of CO_2_. Gravid females displayed elevated pre-CO_2_ baseline flight activity. Male *Ae. aegypti* mosquitoes also showed a strong response to CO_2_ (Figure 1D and 1E).

To study tissue-specific mRNA expression in isolated sensory and neural tissues of male and female *Ae. aegypti*, we dissected individual tissues from pools of non-blood-fed, blood-fed, and gravid female mosquitoes, as well as males. Tissue was flash-frozen and total RNA was extracted and used to generate Illumina RNA-seq libraries from polyA-selected mRNA. From female mosquitoes, we generated libraries from brain, antenna, proboscis, maxillary palp, foreleg, midleg, hindleg, and ovary (Figure 2A). We also dissected the rostrum, a tissue that includes both maxillary palp and proboscis, and the abdominal tip, defined in females as the three terminal abdominal segments, including genitalia and ovipositor (Figure 2A). From male mosquitoes, we generated libraries from brain, antenna, rostrum, foreleg, midleg, hindleg, and abdominal tip, defined in males as the three terminal abdominal segments, including genitalia (Figure 2B). We generated at least three biological replicates for each tissue and subjected each to deep sequencing using an Illumina HiSeq instrument (with the exception of four libraries sequenced on the Illumina Genome Analyzer II) (Figure 2C).

**Figure 2.**
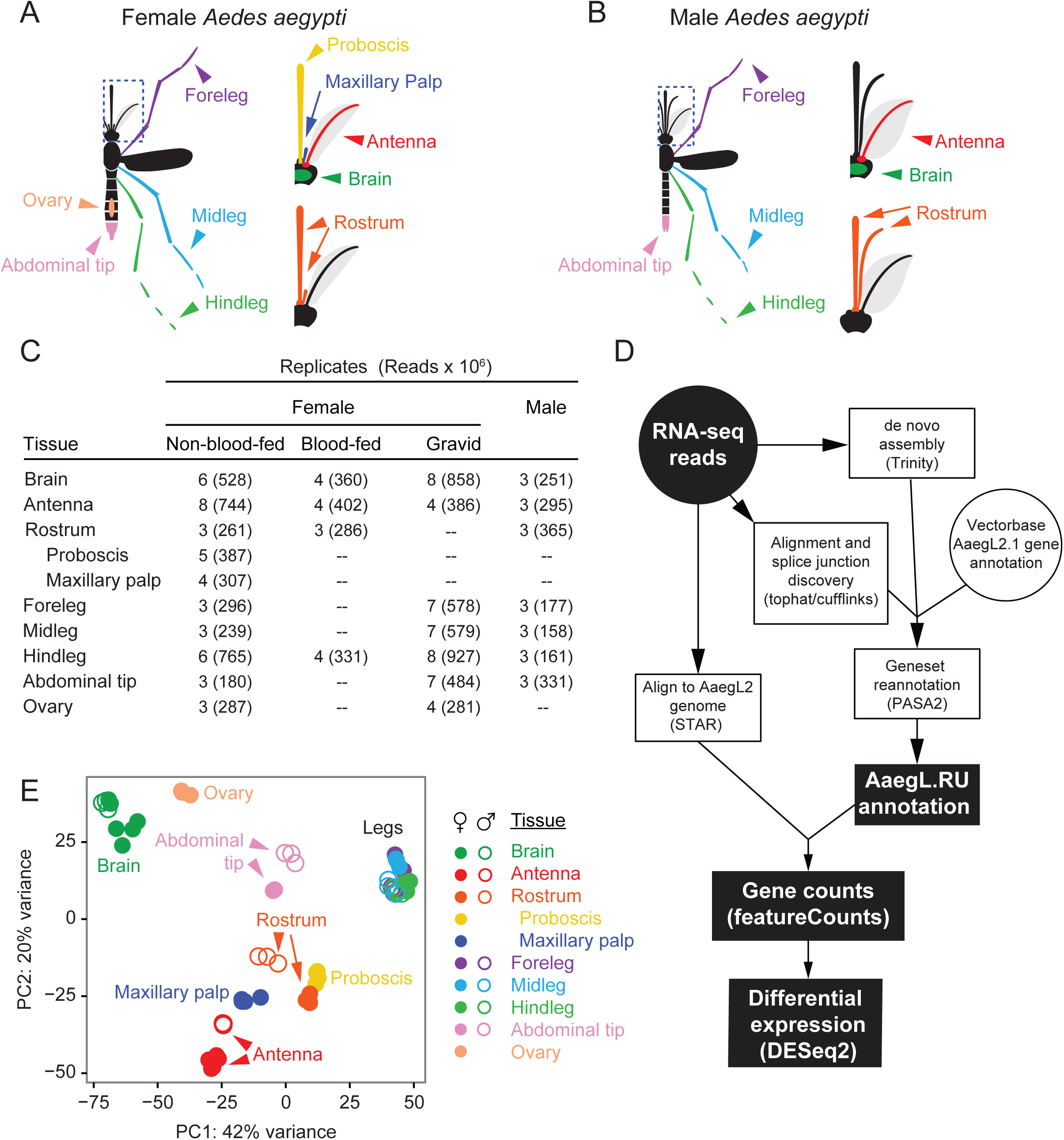
Neurotranscriptome tissues, sequencing, and annotation methods. Cartoon of tissues dissected from adult female (**A**) and male (**B**) *Ae. aegypti*. The rostrum comprises maxillary palp, proboscis, and connecting tissue. Number of biological replicates and sequencing reads across all conditions (**C**) and workflow chart (**D**) of analyses [36, 38–40, 76, 81] used to update the annotation of protein coding genes in the *Ae. aegypti* genome. (**E**) Principal component analysis of transcriptome-wide expression profiles of male and non-blood-fed female tissues.

Correctly quantitating transcript expression from RNA-seq experiments depends on accurate gene models. Before analyzing gene expression across tissues, the two sexes, and the female gonotrophic cycle, we utilized our sequencing data from all tissues from non-blood-fed females and males to update the annotation of protein-coding genes in the *Ae. aegypti* genome (Figure 2D). The depth, replication, and diversity of our sequencing allowed us to re-evaluate the existing annotation of protein-coding genes in the *Ae. aegypti* genome using two complementary approaches: *de novo* transcriptome assembly using Trinity [36] and alignment of sequencing reads directly to the reference genome. By aligning contigs from the *de novo* assembly back to the genome, we were able to combine data generated from these two approaches and use PASA2 software [37] to update existing gene annotations (AaegL2.1; obtained from VectorBase [35]). Reads were also aligned to the genome using STAR [38], and those aligning to genes were counted using featureCounts [39], allowing us to estimate transcript abundance and calculate differential expression at the gene level using DESeq2 [40].

We first carried out a principal component analysis of male and non-blood-fed female libraries to examine the clustering of data by tissue and sex. Large batch effects from library construction methods or problems with tissue contamination during dissection [41] may be revealed by this process. Virtually all of the biological replicates of the same tissue clustered tightly in principal component space, and for brain and legs across the two sexes (Figure 2E).

A comparison of the protein-coding transcriptomes from the community annotations AaegL2.1 and AaegL3.3 and our updated transcriptome, termed AaegL.RU, can be found in Figures 3A–3C. In addition to updating existing gene models, analysis identified 403 putative novel protein-coding genes that did not overlap with existing gene annotations and >25,000 new putative alternatively spliced isoforms as compared to the AaegL3.3 geneset (Figure 3A-B). Furthermore, we identified 169 predicted protein-coding transcripts with orthology to other insect transcriptomes on unaligned contigs from our *de novo* assembly that likely derive from unsequenced portions of the *Ae. aegypti* genome (Figure 3B and Additional file 1). Our data also allowed for the extension of existing gene models, substantially increasing the length of UTRs and coding sequence for many existing genes as compared to the community reference annotations (Figures 3B-C). Together, we propose that there are potentially 572 protein-coding genes missing from the currently published annotation, although we note that our data must be integrated with and evaluated in the context of the large body of other genomic, transcriptomic, and bioinformatics evidence available for *Ae. aegypti* [35]. The results of our reannotation, including novel transcripts, can be downloaded as Additional file 2.

**Figure 3.**
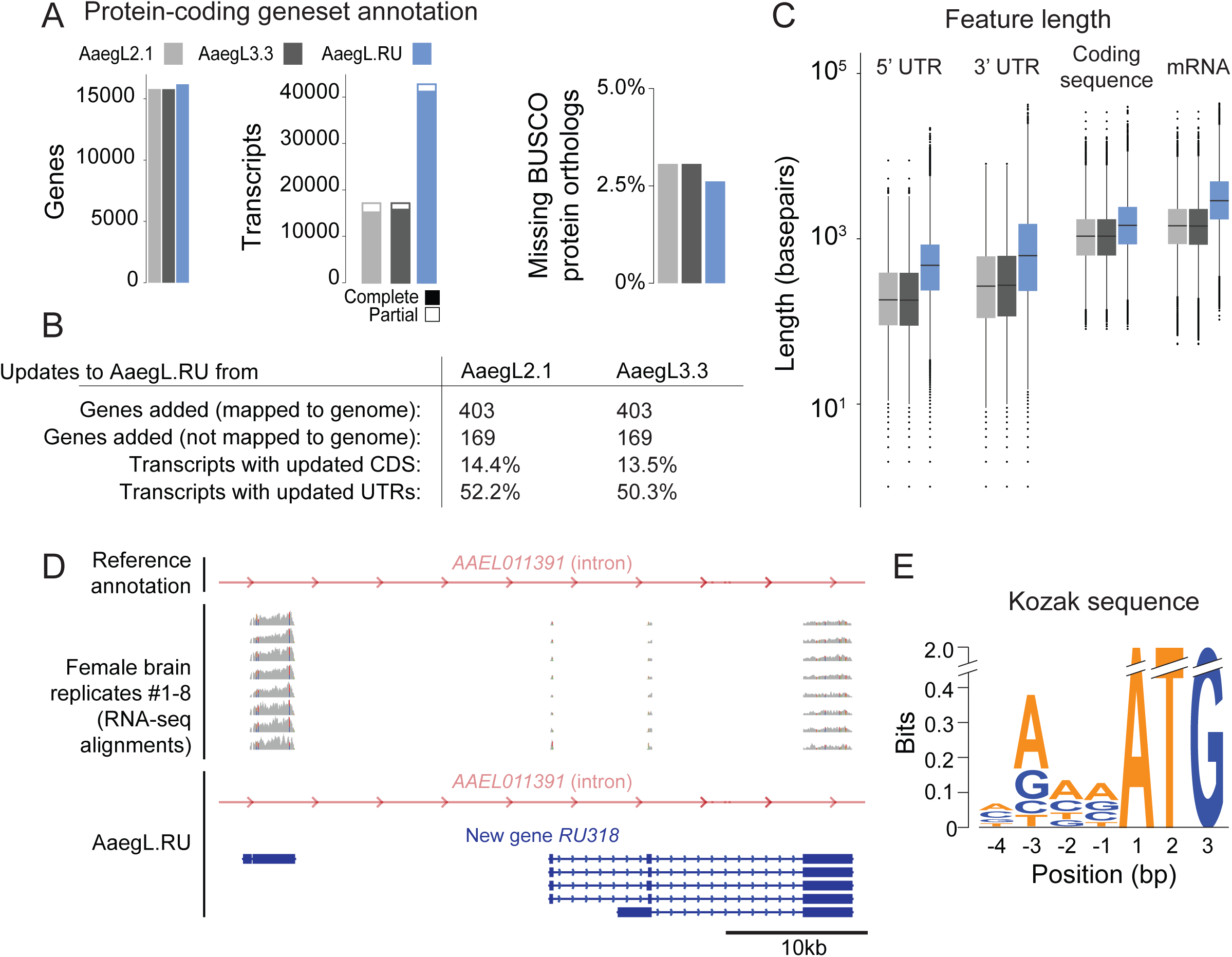
Reannotation of the Ae. aegypti genome. (**A**) Annotation statistics comparing the base annotations (AaegL2.1 and AaegL3.3) with the updated AaegL.RU annotation. (**B**) Summary of added genes and novel predicted genomic loci and transcripts with updated coding sequences as determined by parseva. (**C**) Statistics of all predicted protein-coding transcripts, with data represented as box plots. Median indicated by black line, bounds of box represent first and third quartile, whiskers are 1.5 times the inter-quartile range and outliers denoted by dots. (**D**) RNA-seq evidence from 8 biological replicates supporting the existence of a PASA2-generated gene model for novel gene RU318 within an intron of existing gene model AAEL011391 on supercontig 1.575. (**E**) Sequence logo of the 4 bases upstream of the ATG start codon for all open reading frames from AaegL.RU, comprising the Kozak sequence.

Using our updated geneset annotation, we next classified genes into families related to neuronal function by both incorporating previously published classifications as well as considering their relationship to genes in the well-annotated and -studied vinegar fly *Drosophila melanogaster*. For pre-existing genes, we identified their closest orthologue in *D. melanogaster* using pre-calculated orthology calls of OrthoDB [42]. To account for genes added in our geneset re-annotation, and thus not considered by the OrthoDB databases, we additionally performed blastx of the predicted coding sequence of all transcripts against the *D. melanogaster* proteome (Flybase release 6.06) and report the top BLAST hits with e-values below 0.01 (Additional file 3). Of note, 163 of our 572 proposed novel genes (28.5%) have blastx hits that meet this criterion, as compared to 85.5% of all other annotated genes.

A single example of a novel protein-coding gene, *RU318*, is depicted in Figure 3D. It has high sequence conservation to the *D. melanogaster* TRP channel *waterwitch* [43]. Notably, the current *Ae. aegypti* geneset annotation lacks a predicted orthologue to *waterwitch*, while both other mosquito genomes (*An. gambiae and Cu. quinquefasciatus)* contain orthologues. Based on this sequence similarity, we have included *RU318* in our revised annotation of TRP channels in *Ae. aegypti*. Finally, to aid in transgenesis and other genome engineering approaches, we used all predicted coding sequences to generate a consensus Kozak sequence for *Ae. aegypti* (Figure 3E).

To describe transcript abundance across tissues, we mapped reads from our tissue-specific RNA-seq libraries to the AaegL3 genome and report transcript abundances in units of transcripts per million (TPM) [44]. Mapping statistics for each library can be found in Additional file 4, and TPM values for each replicate library can be found in Additional files 5 and 6. We first described the expression of genes related to neuronal function in specific tissues of non-blood-fed female and male *Ae. aegypti*.

Insects use three major neurotransmitters: acetylcholine is the primary excitatory neurotransmitter in the central nervous system, glutamate is used for neuromuscular transmission from motor neurons to muscles, and GABA is generally considered to be the primary inhibitory neurotransmitter in insect central synapses. Additionally, a number of biogenic amines including serotonin, dopamine, tyramine, octopamine, and histamine function as neurotransmitters or neuromodulators.

We identified acetylcholine receptor subunits by orthology to *D. melanogaster* and *An. gambiae* [45]. Expression of most acetylcholine receptor subunits was highest in male and female brain (Figure 4A). We profiled the expression of three types of glutamate receptors: Kainate, NMDA, and AMPA (Figure 4B). Overall, we found generally broader expression in peripheral tissues as well as brain, consistent with a role in neuromuscular transmission. Ionotropic GABA-A receptors and metabotropic GABA-B receptors were uniformly expressed broadly, with peaks in central brain (Figure 4C). We found two histamine receptor orthologues: HisCl1 and HisCL2 (Figure 4D). Both were expressed in brain at high levels, with HisCl1 (but not HisCl2) also expressed broadly in peripheral tissues. Interestingly, we were unable to identify an orthologue of the glycine receptor *Grd* in *Ae. aegypti* (Additional file 3).

**Figure 4.**
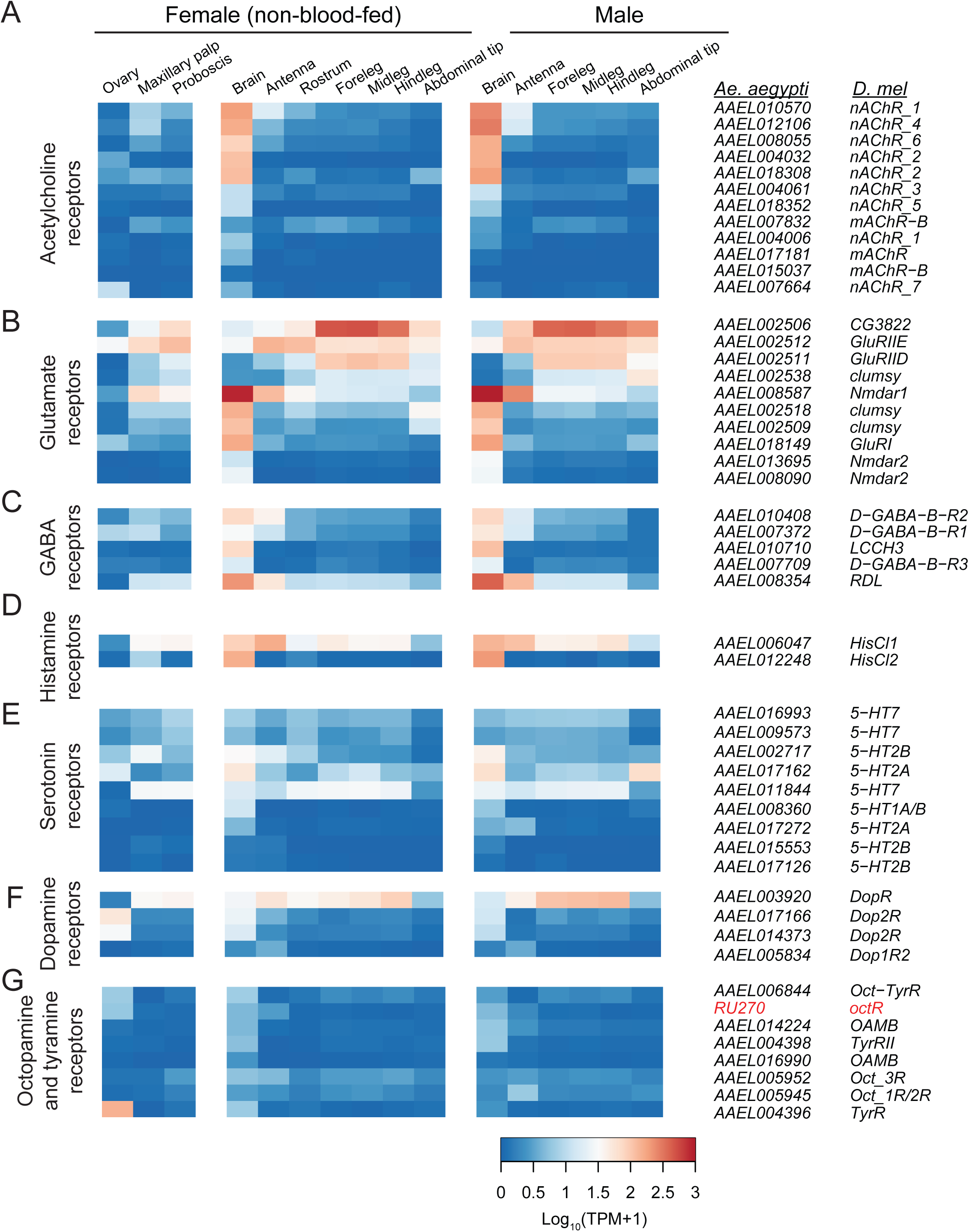
Expression of neurotransmitter-related genes. Expression of receptors for the neurotransmitters acetylcholine (**A**), glutamate (**B**), GABA (**C**), histamine (**D**), serotonin (**E**), dopamine (**F**), and octopamine/tyramine (**G**) in tissues from non-blood-fed female and male mosquitoes. Data are presented as the mean Log_10_(TPM + 1) for all biological replicates from each tissue. AaegL.RU gene set names and the closest orthologue from *D. melanogaster* are indicated on the right.

Biogenic amines represent an important class of neurotransmitters and neuromodulators in insects that have been implicated in processes as diverse as reward, aggression, oviposition choice, and the control of context-specific social behavior [46–49]. *Ae. albopictus* mosquitoes fed constitutively with L-DOPA exhibit lower levels of host-seeking behavior [50]. Serotonin neurons innervate the antennal lobe of *Ae. aegypti* and *An. gambiae* [51], as well as the gut of *Ae. aegypti* [52] and serotonin has been shown to modulate feeding behavior in larval *D. melanogaster* [53]. Most serotonin receptors were expressed at appreciable levels in brain and in various peripheral tissues (Figure 4E). Dopamine receptors had generally variable expression across tissues, including brain, legs, and antennae (Figure 4F). Octopamine/tyramine receptors had generally lower expression values than other receptors, but were detected in brain as well as peripheral tissues (Figure 4G). The *TyrR* orthologue *AAEL004396* was highly and selectively expressed in ovary. We generally observed little obvious sexual dimorphism in neurotransmitter receptor expression. This suggests that there is a gross conservation of neuronal cell types and signaling pathways, at the transcriptional level, across male and female tissues.

Neurotransmitter processing enzymes can serve as cell-type specific markers to reveal the major neurotransmitters produced in particular tissues. The orthologue of *D. melanogaster* glutamic acid decarboxylase 1, or *Gad2*, was expressed in brain and all peripheral sensory tissues, but not ovary. In contrast Gad1, was largely restricted to brain, as well as male abdominal tip (Figure 5A). All dopamine processing enzymes were expressed in male and female brain, including the family of dopamine decarboxylase (Ddc) genes, as well as tyrosine hydroxylase (Th), tyrosine decarboxylase (Tdc), and Tyramine β hydroxylase (Tbh). We also observed strong peripheral expression of Dth and the two Ddc orthologues (Figure 5A). Processing enzymes associated with GABA, glutamate, and acetylcholine were expressed in male and female brain, with many also expressed broadly across the majority of tissues sampled (Figure 5A).

**Figure 5.**
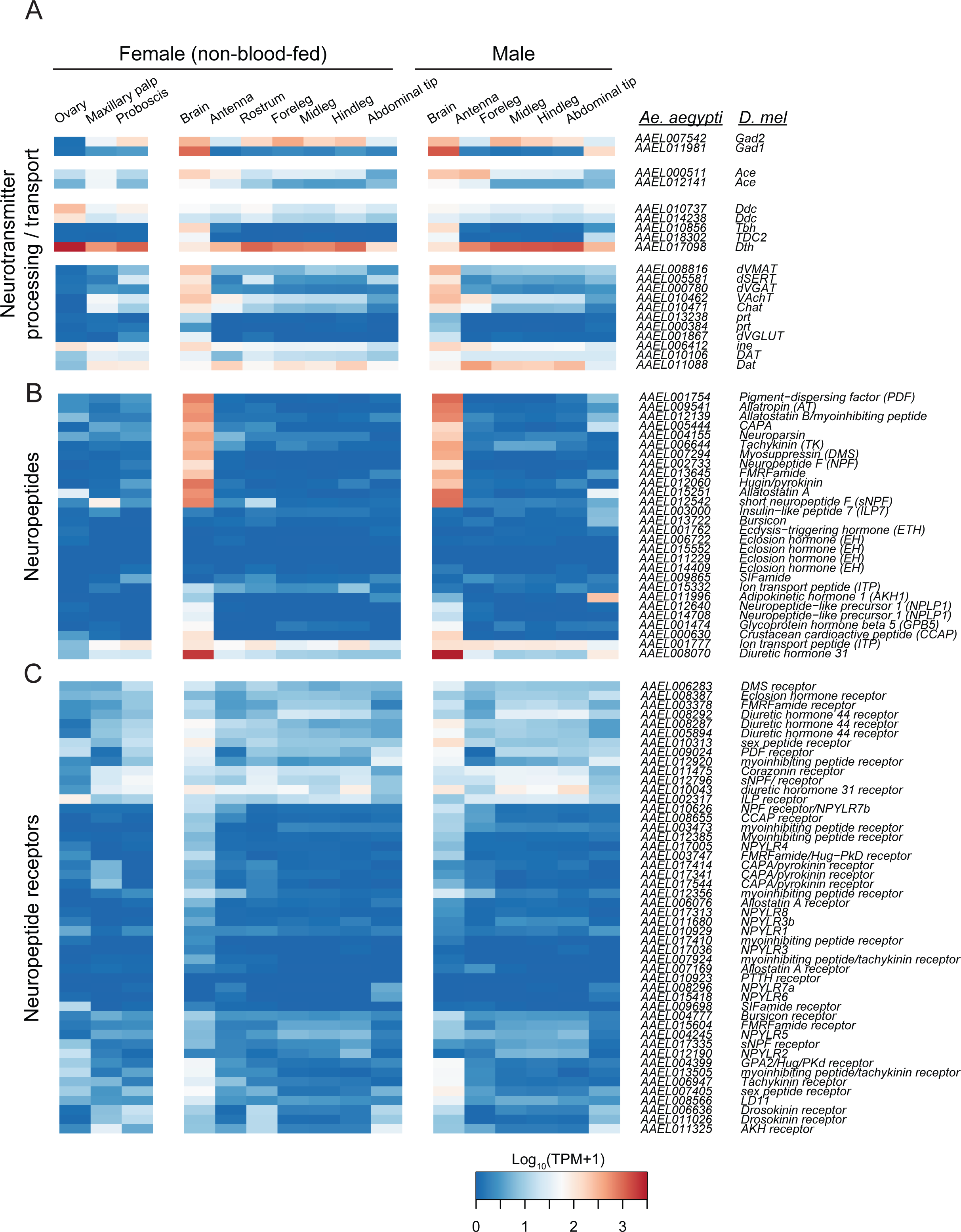
Expression of biogenic amine-related genes. Expression of neurotransmitter transport and processing enzymes (**A**), neuropeptides (**B**), and neuropeptide receptors (**C**) in tissues from non-blood-fed female and male mosquitoes. Data are presented as the mean Log_10_ (TPM + 1) for all biological replicates from each tissue. AaegL.RU gene set names and the closest orthologue from *D. melanogaster* are indicated on the right.

Genes corresponding to *Ae. aegypti* neuropeptides and neuropeptide receptors were defined by orthology to canonical insect neuropeptides and receptors [12, 54, 55]. Many neuropeptides were expressed primarily in brain of male and female mosquitoes, while a number had broader expression patterns that included but were not limited to brain (Figure 5B). This is consistent with reports of the direct detection of neuropeptides in specific regions of the brain, including the antennal lobe [56]. Other peptides, including several predicted orthologues of eclosion hormone (EH), ecdysin-triggering hormone (ETH), and bursicon, were not detected at appreciable levels. We speculate that these genes are expressed selectively in earlier developmental stages that were not sampled in the present study of adult tissues. Neuropeptide receptors had generally broad expression patterns (Figure 5C), indicating that neuropeptides may be centrally produced while exerting anatomically far-reaching humoral effects.

Insects sense chemical substances such as tastants, odorants, pheromones, noxious chemicals, and CO_2_ with an array of chemosensory receptors encoded by large gene families of odorant receptors (ORs), odorant binding proteins (OBPs), ionotropic receptors (IRs), gustatory receptors (GRs), and PPK and TRP ion channels. Other groups have previously used RNA-seq to profile gene expression of a number of these genes in various tissues from several mosquito species [27, 28, 30–34]. Our work builds on these efforts by simultaneously profiling expression of all of these chemosensory genes in multiple individual tissues in *Ae. aegypti* in the same study (Figures 6 and 7).

**Figure 6.**
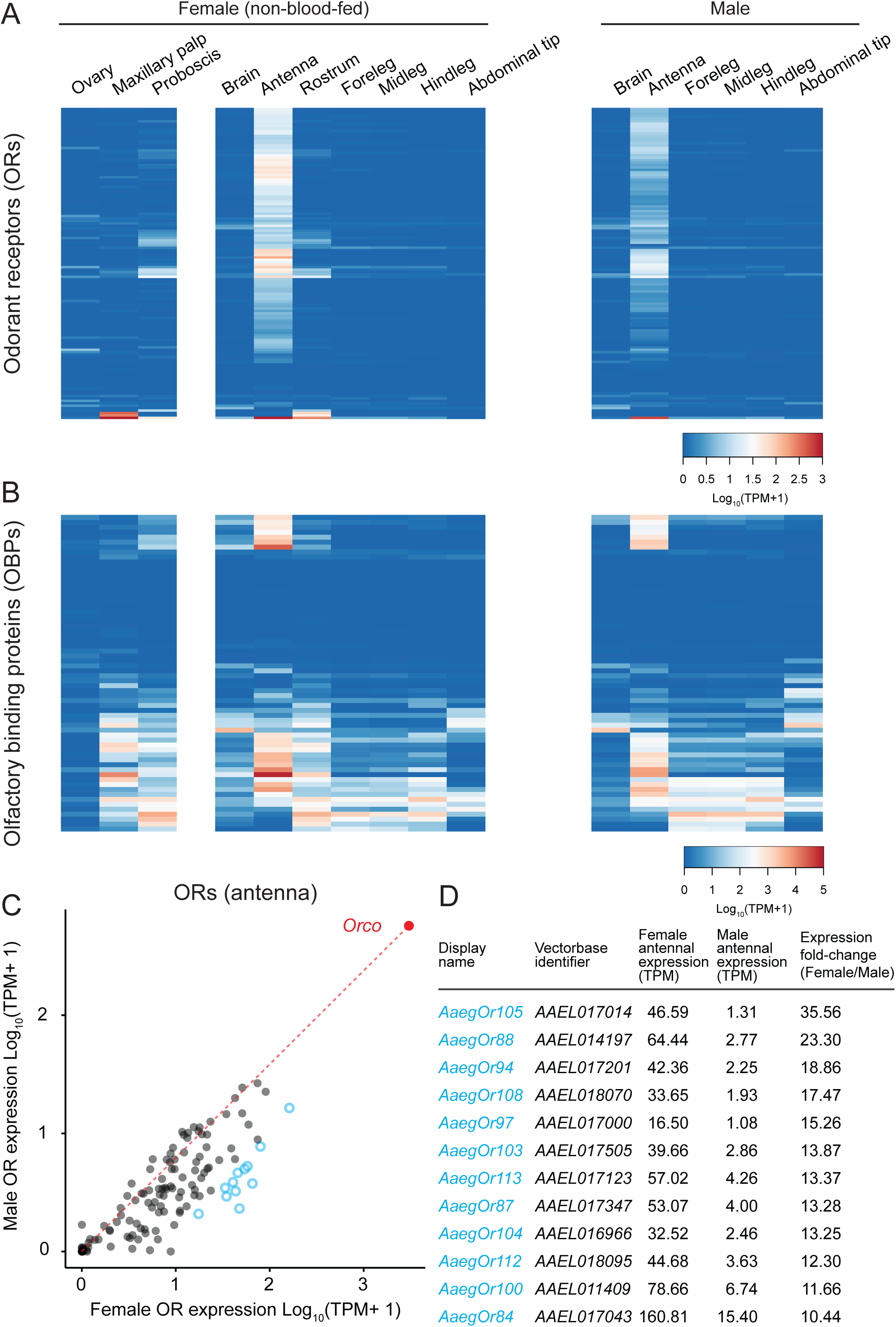
Expression of odorant receptor and odorant binding protein genes. Expression of odorant receptors (ORs) (**A**) and odorant binding proteins (OBPs) (**B**) in tissues from non-blood-fed female and male mosquitoes. Data are presented as the mean Log_10_ (TPM + 1) for all biological replicates from each tissue. Note the expanded scale in panel B. Geneset names and *D. melanogaster* orthologues are omitted because the large number of genes precludes legibility. Raw TPM values for these and all other genes can be found in Additional File 5. (**C**) Scatter plot comparing OR Log_10_ (TPM+1) values in the antenna of non-blood-fed female and male mosquitoes. A line is drawn from the origin to the expression value of the olfactory co-receptor *orco*. Genes in D are represented as cyan circles. (**D**) Table of female-enriched odorant receptors with greater than 10-fold-change in non-blood-fed female versus male antenna and TPM in female greater than 10.

**Figure 7.**
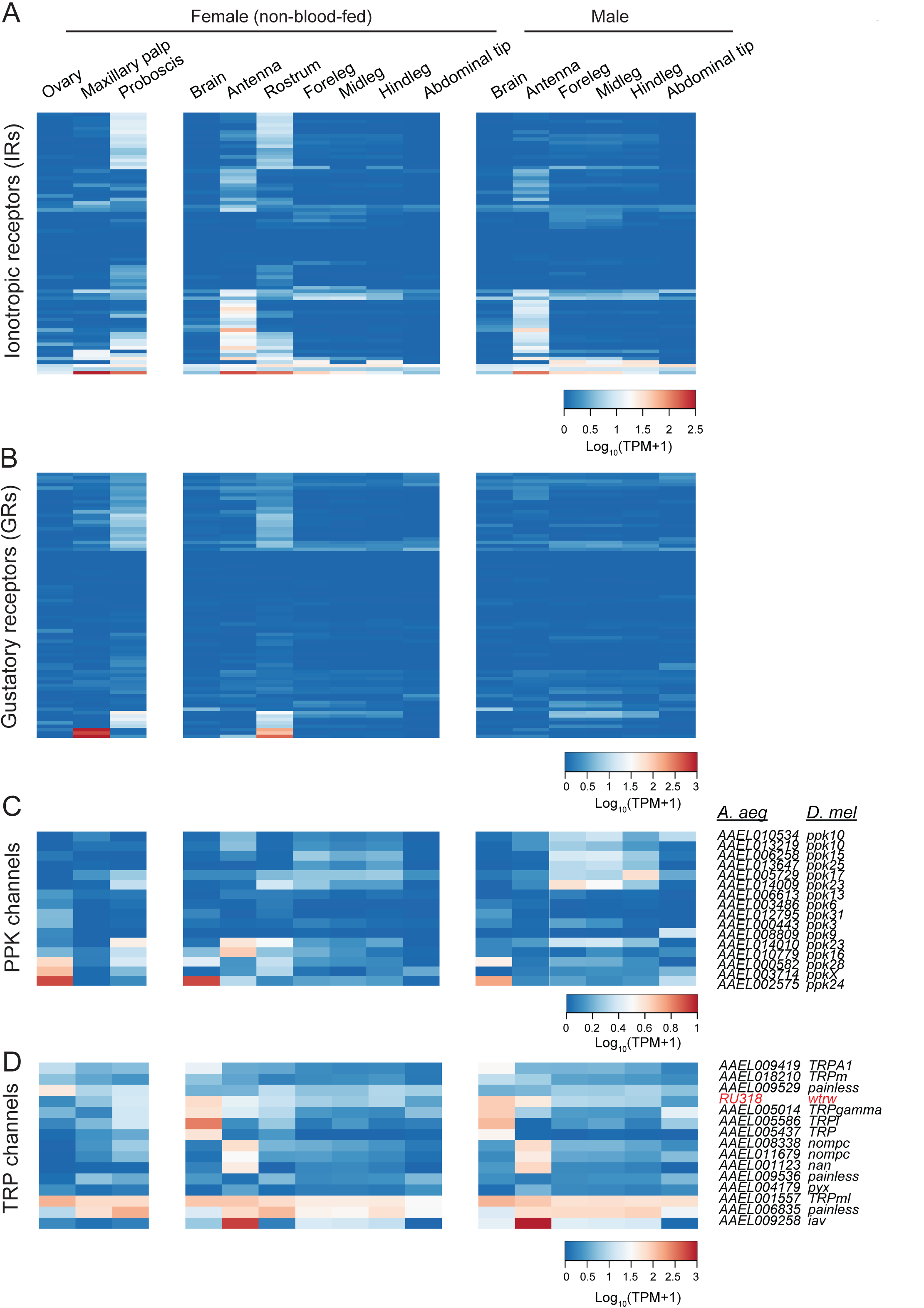
Expression of other chemosensory genes. Expression of ionotropic receptors (IRs) (**A**), gustatory receptors (GRs) (**B**), PPK channels (**C**), and TRP channels (**D**) in tissues from non-blood-fed female and male mosquitoes. Data are presented as the mean Log_10_ (TPM + 1) for all biological replicates from each tissue. AaegL.RU gene set names and the closest orthologue from *D. melanogaster* are indicated on the right, except for IRs and GRs, whose large gene number precludes legibility.

ORs are an insect-specific family of divergent seven transmembrane domain chemoreceptors that sense volatile odors, including pheromones [57]. The majority of ORs are expressed in the antenna, with restricted subsets expressed in either proboscis or maxillary palp (Figure 6A), consistent with previous reports of OR-expressing sensory neurons in these tissues in *An. gambiae* [58]. In contrast to the ORs, OBPs are expressed widely in the tissues profiled here, and vary greatly in their transcript abundance (Figure 6B; note expanded TPM scale relative to Figure 6A). Similar results were found in an analysis of OBP expression in *An. gambiae* mosquitoes [31] and ants [59].

Expression (TPM) values for ORs were broadly elevated in female as compared to male antenna. Antennae of male *Ae. aegypti* are specialized for audition and contain an exaggerated pedicel at their base when compared to female antenna [60]. We speculate that extra cell numbers associated with this enlarged pedicel would effectively dilute mRNA coming from other cells in male antenna, thus reducing the tissue-wide abundance of odorant receptors and other genes expressed in olfactory sensory neurons. To account for these putative differences, we plotted the expression of ORs in male and female antenna (Figure 6C) normalized to the olfactory co-receptor *orco* to reflect the approximate number of olfactory sensory neurons. This normalization depends on the assumption that *orco* expression is not sexually dimorphic, and therefore a reasonable proxy for olfactory sensory neuron number across sexes. Even after accounting for this normalization, we identified 12 OR genes with apparent enrichment in female antenna, with 10- to 35-fold increase in raw expression values as compared to male antennae (Figure 6C and 6D). Because a number of untested assumptions were the basis of these conclusions, we note that these results would need to be validated independently, perhaps by RNA *in situ* hybridization, to gain cellular resolution of gene expression. Increases in mRNA expression could arise either by selective upregulation of gene expression in females, or through developmental changes that would lead to an increase in the number of neurons expressing these receptors in females. Because female mosquitoes show sex-specific chemosensory behavioral responses to odors associated with hosts and oviposition sites, it is not unreasonable to expect sex-specific differences in ORs tuned to these specific odors.

IRs are ligand-gated ion channels derived from variant ionotropic glutamate receptors that tend to be gated by ligands such as acids, aldehydes, and amines [61]. The IR family of *Ae. aegypti* has been previously described [62], and we identified predicted gene models for 2 additional IRs (*RU164* and *RU199*) using blastx against the *D. melanogaster* proteome [Additional files 2-3]. We found three patterns of IR gene expression: those generally restricted to antenna; others selectively expressed in proboscis, rostrum, and maxillary palp; and a small number of IRs expressed across many different tissues examined (Figure 7A). Similar results were previously reported in *D. melanogaster*, *Apis mellifera*, *An. gambiae*, and *Culex quinquefasciatus* [30, 31, 62].

GRs are a family of transmembrane receptors distantly related to ORs [57] that mediate detection of pheromones, tastants, CO_2_ [63], and in the case of *D. melanogaster Gr28b* (*Ae. aegypti Gr19*), light and heat [64, 65]. The annotation of the GR family of *Ae. aegypti* was previously described [66]. GRs are predominantly expressed in the rostrum, maxillary palp, and proboscis (Figure 7B), consistent with their primary and conserved role in taste perception [63]. Notable exceptions include *AaegGr1*, *AaegGr2* and, *AaegGr3* (Figure 7B, bottom 3 genes), which encode CO_2_ receptor genes that function in the maxillary palp [8].

Pickpocket (PPK) channels are a family of amiloride-sensitive DEG/ENaC sodium channels that are involved in the transduction of a number of sensory modalities, including mechanosensation, hygrosensation, and pheromone sensing [67]. We first identified *Ae. aegypti* PPK channels by searching for orthologues to previously described PPKs in *D. melanogaster* and *An. gambiae* [67]. Gene expression profiles of the PPK channels reveal broad expression of most genes across several peripheral tissues (Figure 7C), including proboscis and legs, consistent with a role in various forms of contact chemosensation.

Transient receptor potential (TRP) channels have been implicated in diverse sensory modalities, including heat, light, and chemosensation [68]. *Ae. aegypti* TRP channels were identified by conducting orthologue searches against the 13 identified *D. melanogaster* TRP channels [68]. We identified orthologues to all 13, and two additional genes predicted to be orthologues of *D. melanogaster painless*. Expression of TRP channels was generally broad (Figure 7D), with several interesting tissue-specific expression patterns, most notably in brain and antenna.

In addition to profiling specific gene families, we examined transcripts with sexually dimorphic expression (Figure 8, Additional file 7). We compared expression in 6 tissues between non-blood-fed females and males (Figure 8A-F). Differences may arise from sexually dimorphic expression within individual cell-types or differences in tissue-specific cell-type composition between sexes. To identify a set of broadly dimorphic transcripts, we imposed a more conservative threshold of a fold-change greater than 8 in any tissues, and examined them for overlapping dimorphism across different tissues (Figure 8G and H). Expression patterns of genes that were determined to be dimorphic in at least 3 tissues are indicated in gray in the Venn diagrams in Figure 8G and 8H, and displayed as heat maps in Figure 8I and 8J. We note the enrichment of newly annotated RU genes in the male-specific transcripts, including *myo-sex* (*RU529*) [69] and *nix* (*RU468*) [70]. This likely represents the ability of our RNA-seq data to capture transcripts produced from the *Ae. aegypti* male-specific locus on chromosome 1 that have eluded classical genome sequencing, and therefore annotation, due to large repetitive regions.

**Figure 8.**
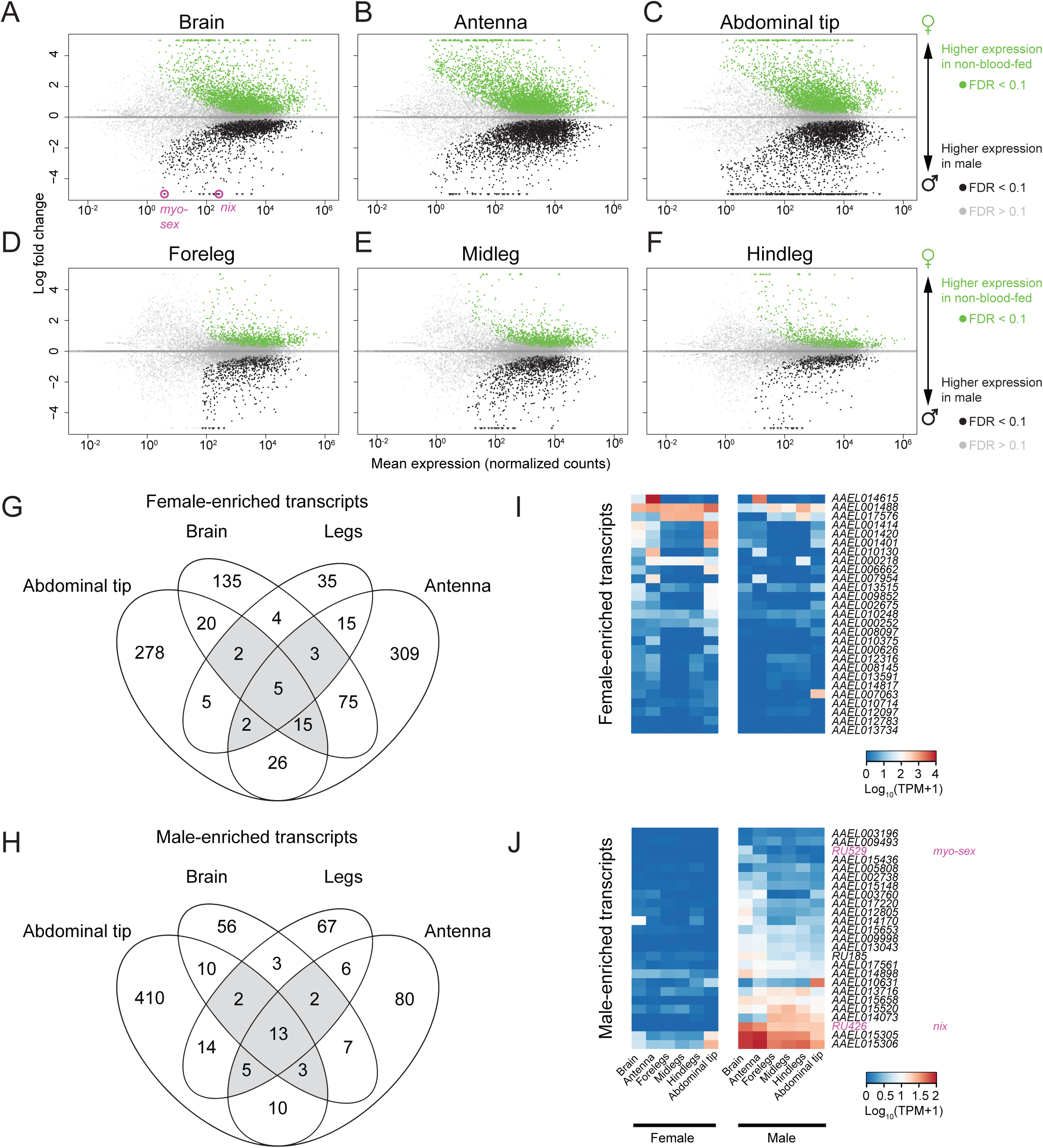
Genes with sexually dimorphic expression. MA plot of differentially expressed genes in brain (**A**), antenna (**B**), abdominal tip (**C**), foreleg (**D**), midleg (**E**), and hindleg (**F**). Genes were identified as significantly regulated within a single tissue using DESeq2 (α < 0.1). Venn diagram of genes that are dimorphic (with a fold-change of at least 8 and an FDR of α < 0.1) in abdominal tip, brain, antenna, or any of the three leg tissues in female (**G**) or male (**H**). Transcript abundance of genes, that are female-enriched (**I**) or male-enriched (**J**) across 3 or more tissues (grey areas of Venn diagrams), sorted by mean abundance in females in descending order (**I**) or males in ascending order (**J**).

Finally, we examined changes in gene expression across the female gonotrophic cycle. After locating and biting a host, female mosquitoes become engorged on a blood-meal that can exceed their unfed body weight. Over the next few days, they must digest this blood and use its nutrients to mature a batch of eggs. During the first 48 hours following a blood-meal, mosquitoes are less responsive to host cues and demonstrate very little locomotion overall (Figure 1B and 1D-1E) [11, 12]. Dramatic changes in gene expression in *An. gambiae* [32, 71], and olfactory function in *An. gambiae* and *Ae. aegypti* [72, 73] after a blood-meal have been documented. Despite these interesting observations, the mechanisms of host-seeking suppression following a blood-meal in *Ae. aegypti* are not well understood. To examine the transcriptional changes associated with this behavioral shift, and begin to approach possible mechanisms, we asked whether there were significant changes in gene expression between non blood-fed, blood-fed, and gravid mosquitoes in brain, antenna, hindleg (Figure 9 and Additional File 8) and several other tissues (Additional Figure S1 and Additional File 8). Genes showing significant changes in expression are displayed in MA plots (Figure 9A–9F). We parsed these data to identify dynamics of gene expression across the gonotrophic cycle in these three tissues. Hundreds of genes showed selective up- and down-regulation in blood-fed relative to non-blood-fed females (Figure 9G, top). Smaller numbers of genes showed peaks of expression in non-blood-fed animals, and suppression in blood-fed and gravid stages, mirroring the behavioral suppression observed in Figure 1. We also identified sets of genes that showed lowest levels of expression at non-blood-fed or gravid stages, and those that had low levels of expression that later peaked in gravid animals. Selected examples of genes belonging to several of these classes of gene expression dynamics are shown in Figure 9H–9K, including the neuropeptide *pyrokinin*/*PBAN* (Figure 9H) and 7 ORs (Figure 9I–9K).

**Figure 9.**
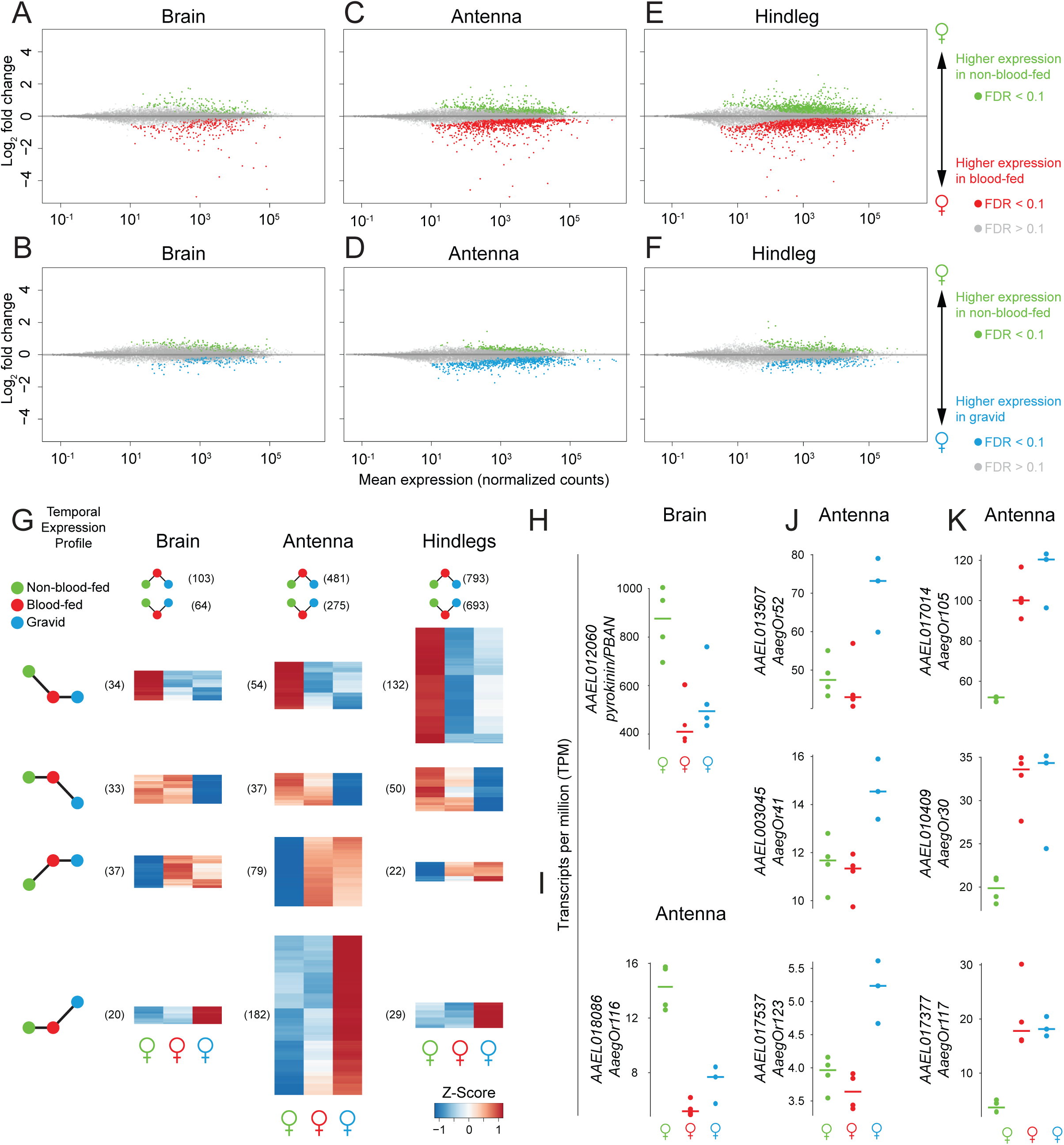
Gene expression changes across the female gonotrophic cycle. MA plots of genes differentially expressed in non-blood-fed versus blood-fed females (**A, C, E**) or non-blood-fed versus gravid females (**B, D, F**) in brain (**A** and **B**), antenna (**C** and **D**), and hindleg (**E** and **F**). Genes were identified as significantly regulated within a single tissue using DESeq2 (α < 0.1). (**G**) Z-scores of significantly regulated genes (α < 0.01) of either non-blood-fed versus blood-fed or non-blood-fed versus gravid) were clustered into 6 qualitatively distinct clusters, four of which are shown as heat maps (**H-K**) Abundances of example genes from brain (H) and antenna (I-K) plotted as TPM.

## Discussion

Here we present the “neurotranscriptome” of brain and peripheral nervous system tissues in female and male *Ae. aegypti* mosquitoes. We used both genome-guided mapping of RNA-seq reads as well as *de novo* transcriptome reconstruction to improve the annotation of existing protein-coding gene models as well as identify 572 putative novel protein-coding genes. By mapping tissue-specific RNA-seq libraries to transcripts generated by these updated gene models, we examined gene expression in 10 female and 6 male tissues from non-blood-fed animals, as well as a subset of tissues from blood-fed and gravid female mosquitoes, representing two important and distinct behavioral states following a human blood-meal.

Given the fragmented state of the *Ae. aegypti* genome and gene annotations, it was important to include a *de novo* assembly approach in our analysis. This allowed us to examine the expression pattern of genes derived from unassembled regions of the genome. For example, a myosin heavy chain gene, *RU529*, is identical in sequence to *Ae. aegypti myo-sex* [69], a gene linked to the sex-determining M-locus of *Ae. aegypti* in a region absent from the current genome assembly. A targeted search for genes in our dataset with similar expression patterns revealed additional genes with male-specific expression. Interestingly, novel unmapped genes from our *de novo* assembly are over-represented in this list and suggest that these may also derive from unassembled genomic loci similar to the M-locus. A recent study confirmed the presence of such a factor, *nix* (identified in our study as RU426), and demonstrated its critical role in sex-determination in *Ae. aegypti* [70].

The present study is a valuable dataset, presenting a comprehensive view of protein-coding gene expression in adult tissues, and yet it remains incomplete. We only sequenced polyadenylated RNA derived from 10 adult tissues. We have not explored the repertoire of small RNAs including microRNAs, non-coding RNAs, or the regulation of alternative splicing. Our re-annotation approach relied on alignments of short reads and *de novo* transcripts back to the current draft of the genome, meaning that gene models residing on misassembled genomic contigs might be incorrectly represented. Indeed, a recent effort to generate a physical map for the *Ae. aegypti* genome found a misassembly rate of approximately 14%, including 6 of the 10 largest supercontigs [23], making it likely that many gene models that rely on the present assembly remain incorrect. Ultimately, a comprehensive annotation of protein-coding genes and non-coding loci within the *Ae. aegypti* genome will require the incorporation of additional genomic sequencing and transcriptomic data derived from distinct developmental stages and tissues [74].

Whole tissue RNA-seq can identify genes differentially expressed across male and female tissues. However, further work will be required to resolve gene expression profiles in individual cells and cell-types. For the purposes of this study, sexually dimorphic transcripts were conservatively defined as those for which the fold-change observed was greater than 8, though we note that there are many more transcripts with less extreme sex-biased expression. Anatomical differences will make it difficult to determine whether observed differences in transcript abundance represent differential regulation within shared cell-types or variation in the cell-type composition of male and female tissues. Interestingly, we describe relatively few examples of sexually dimorphic expression within the chemosensory gene families examined, suggesting that the striking behavioral differences seen between male and female *Ae. aegypti* may be encoded in the neural circuits responsible for the processing of sensory stimuli as opposed to gene expression differences at the sensory periphery.

We do note the statistically significant up- and down-regulation of a handful of olfactory receptors in antenna from gravid females. This is similar to an observed shift in OR expression in the antenna of *An. gambiae* following a blood-meal [32, 71], and suggests that a behavioral shift from host-seeking to oviposition site selection may involve the increased expression of particular ORs tuned to ligands associated with oviposition sites and a concomitant decrease in expression of ORs tuned to host odor. With few exceptions [8, 75], the ligand tuning of specific chemoreceptors has not been determined in *Ae. aegypti.* A systematic effort to de-orphanize *Ae. aegypti* chemoreceptors will be required to address the functional relevance of these observed gene expression changes.

A major goal of this work was to identify gene expression changes correlated with blood-feeding state to gain insight into possible mechanisms by which a blood-meal might influence behavior. We describe many genes that change expression in tissues from blood-fed and gravid mosquitoes, including chemoreceptors, neuropeptides, neuropeptide receptors, and neurotransmitter receptors and processing enzymes, all of which might play important roles in the regulation of behavior and physiology. However, genes from these classes comprise a small minority of all regulated genes, and thus, are unlikely to alone explain the marked shifts in behavior as female mosquitoes transition from host-seeking to oviposition.

We envision this dataset as a resource to guide the selection of candidate genes involved in mosquito behavior as well as providing insight into the principles of gene expression regulation by blood-feeding. Transgenesis of mosquitoes [16] and precisely targeted mutagenesis with tools such as zinc-finger nucleases [3, 8, 12], TALENs [18, 19], homing endonucleases [17], and RNA-guided nucleases [20–22] now allow for the generation of stable mutant lines and other genetic reagents to test the function of candidate genes in mosquito behavior.

## Conclusions

We present a broad view of gene expression in non-blood-fed male and female tissues, focusing particularly on gene families related to neuronal function and chemosensation. We demonstrate that the effects of blood-feeding on gene expression are broad. This study represents the most comprehensive, tissue-specific survey of gene expression in adult *Ae. aegypti* to date and will be foundational in our understanding of the molecular genetic basis of behavior in this important disease vector.

## Materials and methods

### Mosquito rearing

Mosquitoes used in this study were from the genome reference Liverpool strain (LVPIB12) obtained from BEI Resources/CDC/MR-4 (stock number MRA-735). Eggs were hatched in autoclaved water containing ground Tetramin tropical fish food (Tetra) and fed Tetramin food as necessary during larval and pupal development. For routine rearing, adult females were blood-fed on mice under a protocol approved by the Rockefeller University Institutional Animal Care and Use Committee (IACUC Protocol 14756). Male and female adult mosquitoes were reared together under a 14 hr light:10 hr dark cycle under conditions of 25-28°C and 70-80% relative humidity. For female blood-fed libraries, mosquitoes were offered a human arm and allowed to feed to completion. Blood-feeding was verified by separating female mosquitoes with engorged abdomens 24 to 48 hours following blood-feeding. At least 16 hours prior to dissections, mosquitoes were separated under cold-anesthesia into groups of the appropriate size for a given library.

### Mosquito behavior

Uniport experiments were carried out as described [12], with the exception of the stimulus, which was a 11 cm^2^ circle of exposed skin created by cutting a hole in an elbow-length latex glove. CO_2_ concentration in the airstream was measured at 5% with a Carbocap Hand-Held CO_2_ Meter (model GM70, Vaisala Inc.). SciTrackS experiments were carried out as described [8]. Groups of 20 mosquitoes were placed into the flight arena, allowed to acclimate for 15 minutes, and then presented with a 40 second pulse of CO_2_.

### Ethics, consent, and permissions

All blood-feeding procedures and behavioral testing with human subjects were approved and monitored by The Rockefeller University Institutional Review Board (IRB; protocol LVO-0652). Subjects gave their written informed consent to participate in these experiments.

### Tissue dissection and RNA extraction

Mosquitoes were cold-anesthetized and kept on ice until dissections were complete. Individual tissues were removed by forceps or scissors and immediately flash-frozen by placing into nuclease-free tubes in a dry-ice/ethanol bath (-76 °C). The following number of mosquitoes was used for each female library: antenna, 100 to 220; maxillary palp, 126 to 816; proboscis, 275 to 797; rostrum, 110 to 142; brain, 9 to 18; foreleg, 125; midleg, 100 to 125; hindleg, 100 to 138; ovary, 9 to 25; abdominal tip, 50. For male libraries, the following number of mosquitoes was dissected for each tissue library: antenna, 75; rostrum, 40 to 50; brain, 25; foreleg, 100 to 125; midleg, 100 to 125; hindleg, 100 to 125; abdominal tip, 50. Dissected tissue was stored at -80 °C until RNA extraction.

RNA extraction was performed using the Qiagen RNeasy Mini Kit (Qiagen). Tissue was disrupted with an electric tissue grinder loaded with a disposable RNAse free plastic pestle. For legs, abdominal tip, and other cuticle-rich tissue, samples were further disrupted by passing tissue through a QIAshredder Mini spin column (Qiagen). RNA quantity and quality were evaluated using an Agilent BioAnalyzer 2100 and the RNA 6000 Nano Kit (Agilent Technologies).

### RNA-seq library preparation

Libraries from polyA-selected RNA were prepared with TruSeq RNA Sample Preparation Kits (Illumina) or the mRNA sample-prep kit (Illumina), following the manufacturer’s protocol. Between 200 ng and 1 μg of total RNA was used as input for each replicate library. For paired-end libraries, size-selection was performed prior to PCR by gel extraction or by a Pippin Prep instrument (Sage Biosciences) using 2% agarose cassettes containing ethidium bromide. Size selection resulted in libraries with mean insert sizes (excluding sequencing adapters) of 250-450 base pairs (bp). Library quantity and quality were evaluated using an Agilent BioAnalyzer 2100 and the High Sensitivity DNA Kit.

### Sequencing

All sequencing was performed at The Rockefeller University Genomics Resource Center on HiSeq 2000 or Genome Analyzer IIx sequencers (Illumina). All paired end reads were 2 × 101 bp and all single-end reads were 1 × 101 bp with the exception of four 72 bp libraries. Data were de-multiplexed and delivered as fastq files for each library. These sequencing reads are available at the NCBI Sequence Read Archive (SRA) and are associated with BioProject PRJNA236239.

### Transcriptome generation: reference-based mapping

All reads from all libraries were aligned to the AaegL2 reference genome obtained from VectorBase [35] using Cufflinks2, Tophat2, and Bowtie2 software packages [76]. Reads were aligned without respect to existing annotations with the following settings: minimum intron length of 40 bp, maximum intron length of 500 MB. Cufflinks was run on reads from individual conditions and tissues to identify all putative splice junctions, and then combined using *cuffcompare* to identify a consensus set of putative splice junctions identified in our sequencing reads.

### Transcriptome generation: *de novo* assembly

We performed *de novo* assembly as a second approach to reconstruct transcripts from our data. All reads from all libraries were assembled into a genome-free *de novo* assembly using the Trinity software package (version 2013-02-25) [36, 77]. To account for the depth of sequencing, we first performed read normalization to down-sample the number of reads used in the assembly using the included normalization tool in the Trinity software package with a max_coverage setting of 25. Male paired-end, female paired-end, and single-end reads were normalized separately and then combined, resulting in a 49-fold reduction in overall input data. Trinity was run using default settings, with a minimum k-mer coverage of 1, resulting in an assembly with 420,978 contigs.

### Geneset annotation: PASA

The spliced alignments of individual sequencing reads and the alignment of contigs from the *de novo* assembly were used as input to PASA2 [37] as a means of updating the reference gene annotation using the software’s alignment assembly and annotation comparison workflow. Briefly, *de novo* contigs were aligned to the genome (AaegL2; VectorBase [35]) using the short read aligners BLAT and GMAP. These alignments were combined with the combined cufflinks output from genome-guided mapping to create assemblies of spliced alignments. These assemblies were compared to reference annotations (AaegL2.1; VectorBase [35]) and used to extend, update, or merge reference annotations. Additionally, this analysis identified 403 putative protein-coding genes not covered by the current annotation (see “Geneset annotation: naming of genes and geneset comparisons” below). Default PASA2 parameters were used with the exception of the number of allowed exons in 5’ or 3’ UTRs (--MAX_UTR_EXONS=3). Due to a technical oversight, 5 genes were added manually after the PASA2 run, using previously published coordinates: *AaegGr27*, *AaegOr54*, and *AaegIr41d.2*, *AaegIr75k.4*, and *AaegIr7h.2*.

### Identifying novel unmapped genes

To identify novel transcripts that do not map to the current genome assembly, we filtered our *de novo* assembly as follows. First, we excluded all contigs that mapped to the genome or to cDNA from an existing transcript using GMAP. Next, we required that each contig encoded a complete open reading frame (ORF) of at least 30 amino acids in length, as predicted by transdecoder (http://transdecoder.sourceforge.net). We then screened for likely bacterial and fungal contamination by performing blastx with default settings of the remaining contigs against the nr database (NCBI), and excluded anything for which the top hit was fungal, bacterial, or mammalian. Finally, we performed blastx of each remaining contig against a database of insect transcriptomes (*Anopheles gambiae* [AgamP3], *Apis mellifera* [Amel_4.0], *Culex pipiens* (now *Culex quinquefasciatus)* [CpipJ1], *Drosophila pseudoobscura* [r3.1], *Heliconius melpomene* [v1.1], *Ixodes scapularis* [IscaW1.2], *Nasonia vitripennis* [Nvit_1.0], *Rhodnius proxlixus* [RproC1], *Bombyx mori* [SilkDB v1.0], *Drosophila melanogaster* [r5.50] and *Triboleum castaneum* [v2.0; without mitochondria], requiring that there was a match with an e-value of less than 0.01. 232 contigs that passed these conservative filters were considered to be high-confidence novel genes derived from portions of the genome that have not been sequenced or had assembly problems. These 232 contigs were collapsed using CD-HIT [78, 79] resulting in 169 novel transcripts that were included in downstream analysis.

### Geneset annotation: naming of genes and geneset comparisons

To name each gene in our updated geneset, we first compared them to existing annotations in AaegL3.3 using cuffcompare [76] and carried over accession numbers for those loci that were highly similar to existing annotations. Genes that did not match existing loci in these cuffcompare analyses are numbered sequentially as RU1–RU572 (Additional File 2). For genes with a VectorBase accession number, orthology to *Drosophila melanogaster* was retrieved from OrthoDB (ODB8, dipteran dataset) [42]. We used NCBI blastx to compare CDS for all transcripts against the *D. melanogaster* proteome; results for up to 10 hits with an e-value of less than 0.01 are listed in Additional File 3. To determine the proportion of transcripts that were updated as compared to the VectorBase annotations (Figure 3C; AaegL2.1 and AaegL3.3), we used the parseval tool in the AEGeAn package to compare gff3 annotation files [80].

### Library QC

A file detailing library alignment and quantitation statistics is available as Additional file 4 While all sequencing reads were used for genome reannotation, aberrant clustering of transcriptome-wide expression patterns from two non-blood-fed female brain libraries resulted in their exclusion pool and the DESeq2 model was re-run. Additionally, signs of contamination of male rostrum libraries resulted in their removal from expression analysis (note that these are retained in the PCA plot in Figure 2E).

### Expression data and differential expression analysis

All reads from individual libraries were mapped to the AaegL3 genome using STAR version 2.4.1c [38], and reads mapping to each gene in the AaegL.RU or AaegL3.3 geneset annotation were counted at the gene level using featureCounts v1.4.6-p3 [39] (Additional File 3). For abundance visualization, raw counts were converted to TPM [44] in R. Raw counts were used for differential expression analysis in R using DESeq2 v1.8.1 [40], and the PCA analysis in Figure 2E was performed with DESeq2 using counts subjected to Variance Stabilizing Transformation (VST).

Sexually dimorphic genes were identified with a DESeq2 model incorporating all non-blood-fed female and male libraries from a single tissue and visualized as MA plots generated with significance indicated at an FDR of α < 0.1 (Figure 8A-F, Additional File 7). Venn diagrams were generated using the R library VennDiagram (Figures 8G and 8H). Transcript abundance of genes identified as dimorphic in at least three tissue groups were visualized as heat maps sorted by the sum of the TPM in the dominant sex (Figures 8I and 8J).

Genes regulated by blood-feeding state were identified in DESeq2 with single-tissue models incorporating single-end libraries from female tissues (with the exception of ovary, where all libraries were paired-end). In tissues with three time-points (brain, antenna, and hindleg), Z-scores of expression for genes with an FDR of α < 0.01 (in either comparison) were generated using the ‘scale’ function of R, and clustered using the hclust(method = ‘complete’) and dist(‘method = euclidean’) functions in R. (Figure 9G).

## Data availability

All raw reads, gene set annotations, expression data, and sequences of new genes generated under this project have been deposited in the NCBI SRA under BioProject number PRJNA236239.

## Abbreviations

5-HT: 5-hydroxytryptamine (serotonin)
bp: base pair
CDS: coding sequence
CO_2_: carbon dioxide
CV: coefficient of variation
DA: dopamine
Ddc: dopamine decarboxylase
DNA: deoxyribonucleic acid
EH: eclosion hormone
EST: expressed sequence tag
ETH: ecdysin-triggering hormone
GABA: □-aminobutryic acid
IACUC: Institutional Animal Care and Use Committee
IR: ionotropic receptor
IRB: Institutional Review Board
kb: kilobase
m: meter
MA plot: plot using an M (log ratios) and A (mean average) scale
MB: megabase
mL: milliliter
NPY: neuropeptide Y
OBP: odorant binding protein
OR: odorant receptor
ORF: open reading frame
PCA: principal component analysis
PCR: polymerase chain reaction
PPK: pickpocket
RNA: ribonucleic acid
RNA-seq: mRNA-sequencing
sec: seconds
Tbh: Tyramine β hydroxylase
Tdc: tyrosine decarboxylase
Th: tyrosine hydroxylase
TPM: transcripts per million
TRP: transient receptor potential
μL: microliter
UTR: untranslated region
VST: variance stabilizing transformation

## Authors′ contributions

BJM, CSM, and MJD dissected tissues and made libraries and together with LBV conceived and planned the study; BJM carried out the experiments in Figure 1; BJM, CSM, and OD carried out bioinformatic analysis. BJM and LBV wrote the paper and produced the figures, with input from the other authors.

## Description of additional data files

The following additional data are available with the online version of this paper.

**Additional file 1** is a FASTA file with the raw sequences for unmapped genes

**Additional file 2** is a GFF3 file with the complete annotation of all genes in the AaegL.RU geneset

**Additional file 3** is an annotation table for all genes in AaegL.RU, including OrthoDB orthologue groups shared with *D. melanogaster*

**Additional file 4** describes the mapping and read count statistics for each individual RNA-seq library

**Additional file 5** is a table of transcript abundance (in units of TPM) for all non-blood-fed and male tissues, AaegL.RU annotation

**Additional file 6** is a table of transcript abundance (in units of TPM) for all non-blood-fed and male tissues, AaegL3.3 annotation

**Additional file 7** is a spreadsheet describing the DESeq2 differential expression statistics of sexually dimorphic transcripts for all applicable tissues

**Additional file 8** is a spreadsheet containing the DESeq2 differential expression statistics of transcripts regulated by blood-feeding state for all applicable tissues

**Additional Figure 1**: Gene expression changes across the female gonotrophic cycle. MA plots of genes differentially expressed in non-blood-fed versus blood-fed female rostrum (**A**) or non-blood-fed versus gravid female forelegs (**B**), midlegs (**C**), ovary (**D**) and abdominal tip (**E**). Genes were identified as significantly regulated by a single-tissue comparison using DESeq2 (α < 0.1).

## Acknowledgements

We thank Nicolas Robine, and members of the Vosshall Lab for discussions and comments on the manuscript; Felix Baier and Chloe Goldman for expert technical assistance; Katie Kistler for assisting with the behavioral experiments in Figure 1b and c; Deborah Beck and Gloria Gordon for mosquito rearing; Daniel Lawson for discussion and sharing annotations in progress at VectorBase; Connie Zhao and James Hughes for facilitation of DNA sequencing at the Rockefeller University Genomics Resource Center, and Scott Dewell for bioinformatic support. This work was funded in part by a grant to R. Axel and L.B.V. from the Foundation for the National Institutes of Health through the Grand Challenges in Global Health Initiative and the following National Institutes of Health grants: K99 award from NIDCD to CSM (DC012069), an NIAID VectorBase DBP subcontract to LBV (HHSN272200900039C), and a CTSA award from NCATS (5UL1TR000043). BJM was supported by Henry and Marie-Josée Kravis and Jane Coffin Childs Postdoctoral Fellowships. LBV is an investigator of the Howard Hughes Medical Institute.

---

**Supplementary Figure S1.**
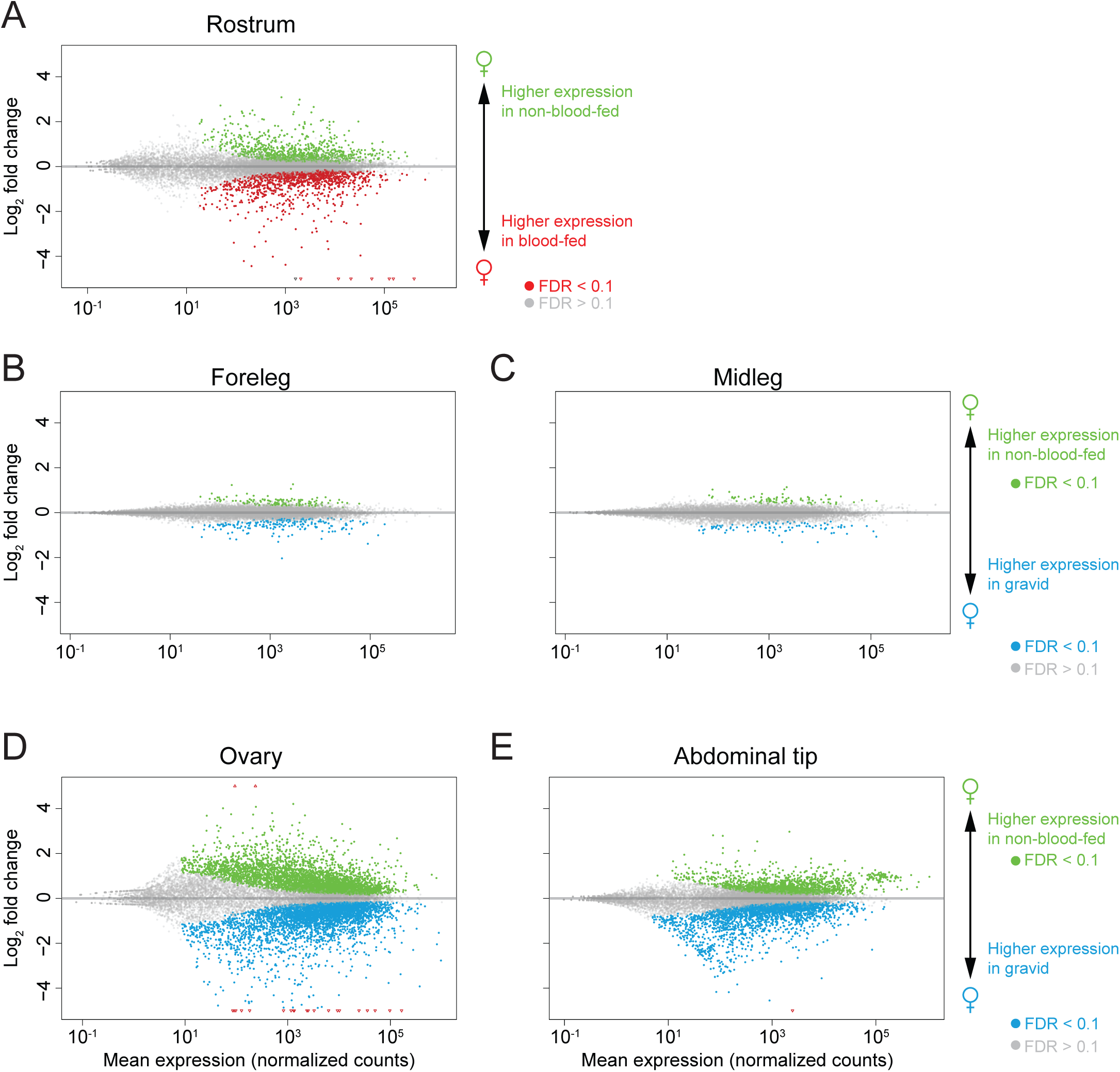
MA plots of genes differentially expressed in non-blood-fed versus blood-fed females (**A**) or non-blood-fed versus gravid females (**B-E**) in rostrum (**A**), foreleg (**B**), midleg (**C**), ovary (**D**), and abdominal tip (**E**). Genes were identified as significantly regulated within a single tissue using DESeq2 (α < 0.1).

